# Integrative multi-omic analysis reveals oral microbiome-metabolome signatures of obesity

**DOI:** 10.1101/2025.06.22.659458

**Authors:** Ahmed A. Shibl, Tsedenia W. Denekew, Anique R. Ahmad, Salah Abdelrazig, Christopher E. Leonor, Lina Utenova, Guihao Zhang, Mamoun AbdelBaqi, Yashaswi Malla, Muhammad Arshad, Marc Arnoux, Nizar Drou, Abdishakur Abdulle, The UAE Healthy Future Study Investigators Group, Raghib Ali, Shady A. Amin, Youssef Idaghdour, Aashish R. Jha

## Abstract

Obesity is a major global health challenge and a leading risk factor for cardiometabolic disorders. The global surge in obesity, driven by industrialization and the widespread consumption of low-fiber, ultra-processed food, highlights an urgent need for deeper biological insights. While the gut microbiome has been studied in the context of obesity, the contribution of the oral microbiome—the second largest microbial ecosystem in the human body–remains largely underexplored. Here, we report findings from a deeply-phenotyped prospective-cohort of 628 Emirati adults, leveraging amplicon sequencing of mouthwash samples and multi-omics profiling and functional and metabolic activity analysis of 97 obese individuals and 95 matched controls, making this the most comprehensive multi-omics analysis of the oral microbiome. We identified significant differences in oral microbial diversity, composition, functional pathways, and metabolic profiles between obese and non-obese groups. Integrated multi-omics analysis of the 192 matched metagenomes and metabolome samples uncovered significant metabolic reprogramming and altered energy regulation in obesity. Specifically, the oral microbiome of obese participants were enriched for the proinflammatory *Streptococcus parasanguinis* and *Actinomyces oris,* and the lactate-producing *Oribacterium sinus.* Many microbial pathways involved in dietary carbohydrate metabolism, histidine degradation, as well as the production of obesogenic biomolecules were also enriched in obese participants; however, B-vitamin and heme production pathways were depleted. Consequently, metabolites resulting from these pathways such as lactate, histidine derivatives, choline, uridine, and uracil were elevated in obesity. These consistent microbiome-metabolite shifts were strongly associated with prominent obesity-associated cardiometabolic markers, including serum triglycerides and alkaline phosphatases, establishing a robust link between oral microbiome and obesity. These findings provide the most comprehensive insights into how disrupted microbial-metabolic cross-talk in the oral cavity may contribute to obesity and related cardiometabolic disease risk, underscoring the potential of targeting of oral microbiome-host interactions as a novel avenue for obesity prevention and intervention.

## Introduction

The World Health Organization reports that nearly half of the global adult population (43%) is overweight, with 890 million individuals aged 18 or older classified as obese^1^. Obesity, characterized by excessive fat accumulation, is a major risk factor for a range of metabolic disorders, including diabetes, hypertension, and cardiovascular diseases^2^. The prevalence of adolescent obesity has surged fourfold in recent decades, representing an escalating global health challenge^1,2^. While genetic predisposition plays a role in weight gain^3^, the worldwide increase in obesity is largely attributed to changing dietary patterns accompanying industrialization, marked by increased consumption of ultra-processed, low-fiber foods^2,3^. This trend is especially evident in countries such as the United Arab Emirates (UAE), where rapid urbanization has catalyzed significant dietary and lifestyle shifts^4^, contributing to marked rise in obesity and cardiovascular mortality in the Emirati population^5^.

Disruption in the gut microbiome is recognized as a key driver of obesity development and progression^6,7^, along with obesity-associated comorbidities^8,9^ by modulating several physiological processes, including carbohydrate metabolism, energy homeostasis, vitamin production, and body weight regulation^10,11^. Although the role of the gut microbiome in obesity and other chronic diseases has been extensively studied^9–11^, the potential contributions of the oral microbiome to systemic health and disease remain underexplored^12^. The oral cavity harbors a diverse array of microorganisms such as bacteria, archaea, viruses, and unicellular eukaryotes, which produce a variety of biomolecules^13^. This collective community of microorganisms and their products constitute the oral microbiome—the second largest microbial ecosystem in the human body after the gut^14^. Compounds generated by oral microbes can interact with the epithelial lining of the oral cavity, triggering various signaling mechanisms^13^. Additionally, these biomolecules can also enter the bloodstream by crossing the well vascularized buccal mucosa and interact with distant host tissues. Lifestyle and dietary factors, such as high sugar intake, processed food consumption, smoking, poor oral health, and antibiotics use can significantly affect the oral microbiome integrity^15,16^.

Emerging evidence links dysbiosis of the oral microbiome to several metabolic diseases, including obesity^17^. Obese individuals often show decreased overall microbial diversity, along with increased relative abundances of Bacillota (previously known as Firmicutes), periodontal pathogens, and pro-inflammatory bacteria in their oral microbiomes^17,18^. These microbial shifts may contribute to systemic inflammation and metabolic dysfunction^19^. Furthermore, obesity related shifts in oral microbial community composition^20^ may induce alterations in the gut microbiome and collectively they can influence systemic changes in metabolic diseases^21^. Also, the practicality of sampling the oral microbiome makes it particularly appealing for large-scale and longitudinal studies. However, the insights gained from oral microbiome studies have been constrained by small sample sizes and reliance on 16S ribosomal RNA (16S rRNA) gene amplicon sequencing^22^. Whole metagenomics shotgun sequencing offers a more robust method for identifying disease-associated species and their functions, whereas metabolomics detects shifts in oral metabolites that act as messengers, conveying information between the mouth and the host^13,22^. Integrating metagenomics and metabolomics provides a promising strategy for gaining a comprehensive understanding of how oral microbes contribute to diseases^23^. Here, we implement multiple methodologies to investigate the role of oral microbiome in obesity in a large cohort. First, we utilize 16S rRNA sequencing in a large discovery cohort comprising 628 individuals to identify various lifestyle and health associated factors contributing to oral microbiome variation, which led us to identify obesity as a significant contributor. Next, we conduct an in-depth obesity-focused multi-omics association analysis using shotgun metagenomics and untargeted metabolic profiling by liquid chromatography followed by mass spectrometry (LC-MS) on mouthwash samples from 192 individuals (97 obese and 95 matched healthy-weight controls). This makes our study the most compreshensive multi-omics analysis of the oral microbiome to date. Integrating these diverse datasets using a multi-omics framework allows us to identify distinct oral microbial species, bacterial enzymes, pathways, and metabolites altered in obesity. Finally, we successfully correlate these oral microbial and metabolic alterations with key obesity-associated clinical biomarkers derived from blood and urine samples, establishing a robust link between oral microbiome and obesity. These findings offer initial insights into how compositional, functional, and metabolic shifts in the oral microbiome may contribute to obesity.

## Results

### Study design and cohort description

The UAE Healthy Futures Study (UAEHFS) is a large, ongoing population-based prospective cohort study of 20,000 Emirati nationals aimed at identifying factors contributing to cardiometabolic diseases of significant public health concern^24^. In this study, we included a cohort of 669 consenting Emirati nationals aged 18-43 years (mean±SD= 24 ± 5.3 years) enrolled in the UAEHFS, from whom mouthwash samples were collected. Participants completed standardized survey questionnaires capturing demographic (age, sex, marital status, etc.) and lifestyle (smoking, exercise, etc.) attributes, and underwent clinical assessments for anthropometric and physiological traits (BMI, blood pressure, heart rate, etc.). Based on self reported questionnaires, 13% of participants reported a prior history of periodontal disease. However, ∼95% brushed at least once daily and reported no current symptoms of poor overall health (e.g. mouth ulcers, loose teeth, or painful gums), suggesting that most participants had generally healthy oral conditions at the time of sampling. Additionally, we measured 58 clinical parameters from blood and urine (**Fig. 1a**). The average BMI and cholesterol levels in this cohort were 26.8 kg/m^2^ (SD= 6, range= 14.4-52.2 kg/m^2^) and 179 mg/dL (SD= 38.3 mg/dL, range= 93-445 mg/dL), respectively. Consistent with previous reports on the broader Emirati population^5,25^, obesity (BMI ≥ 30, 30.4%) and hypercholesterolemia (blood cholesterol ≥ 200 mg/dL, 28.4%) were the two most prevalent medical conditions in this cohort.

**Figure 1.**
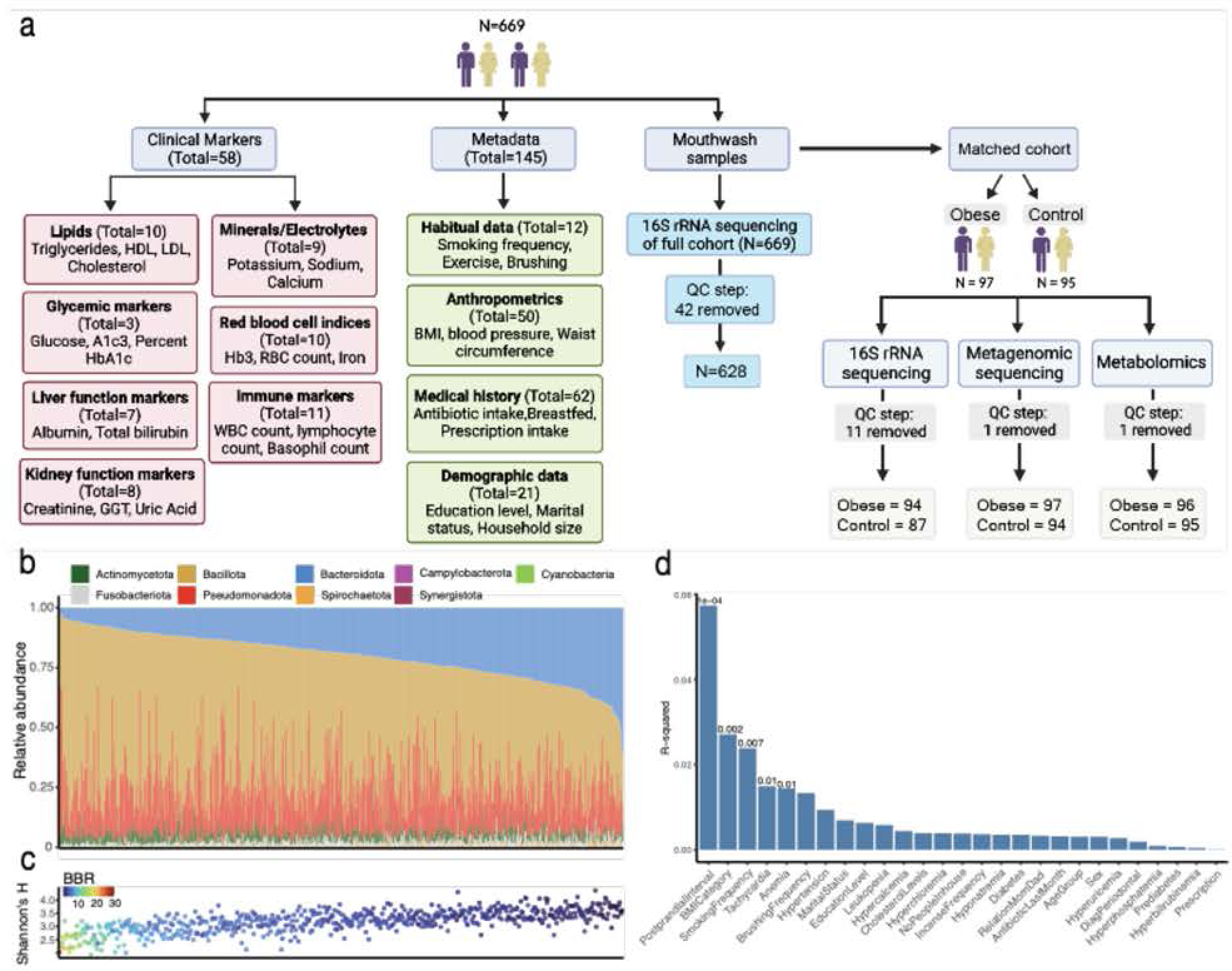
Study design, cardiometabolic conditions, and oral bacteria in the full cohort. **a,** A total of 669 mouthwash samples were collected from the Emirati nationals enrolled in the ongoing UAEHFS for 16S rRNA gene sequencing (blue boxes). A total of 58 clinical biomarkers from blood and urine samples were assessed from these individuals (pink boxes) and a total of 145 demographic, lifestyle, and anthropometric variables were collected from self-completed questionnaires or physical exams (green boxes). In addition, a subset of 192 individuals that included 97 obese and 95 matched healthy-weight individuals (controls) were used for shotgun metagenomics sequencing and untargeted LC-MS metabolomics. **b,** Relative abundances of the bacterial phyla in the mouthwash samples in 628 participants retained after quality control shows the interindividual variation in oral microbiota. **c,** Alpha diversity calculated using Shannon’s Diversity Index is strongly correlated with the Bacillota-to-Bacteroidota ratio (BBR). For **b** and **c**, participants are sorted by the relative abundance of Bacteroidota on the x-axis. **d,** Postprandial-interval, BMI categories, smoking frequency were most strongly associated with oral microbiome composition. Tachycardia and anemia were also associated but their effect sizes were relatively small.

### Effect of lifestyle and systemic health on the oral microbiome in the Emirati population

The diverse community of microbes residing in the oral cavity plays a crucial role in oral health^13,26^. However, the link between variations in oral microbiome and broader lifestyle and systemic health factors beyond the oral cavity remains largely unexplored^22^. Identifying lifestyle and health-related variables contributing to the oral microbiome is essential before conducting association studies, as these factors may act as confounders. To address this, we initially characterized the oral microbiome of 669 participants using 16S rRNA amplicon sequencing, from which 628 passed our quality control criteria. Consistent with previous studies^13^, Bacillota (previously known as Firmicutes, 14.9-87.3%), Bacteroidota (previously known as Bacteroidetes, 2.7-61.5%), and Pseudomonadota (previously known as Proteobacteria, 0.09-63.6%) were the predominant phyla, collectively accounting for ∼93% of reads across the 628 samples (**Fig. 1b**). Oral microbial diversity, calculated using Shannon’s Diversity Index, was significantly associated with postprandial interval (*P*=4×10^−5^, *GLM*). Moreover, it was negatively correlated with the Bacillota-to-Bacteroidota ratio (BBR) (**Fig. 1c**, *Spearman’s ρ*= −0.71, *P*<2.2×10^−16^, *Spearman’s Rank Correlation Test*). Notably, we did not find consistent associations between oral microbial diversity and other anthropometric, demographic, lifestyle, or health factors examined.

A Principal Coordinate Analysis (PCoA) of the amplicon sequence variants (ASVs) from these participants, performed using weighted UniFrac distances, revealed that oral bacterial composition was significantly associated with BMI categories (*P*=0.018, *EnvFit*) along with other lifestyle factors such as postprandial interval and smoking. We assessed the relationship between oral microbiome composition and 27 non-collinear metadata variables—13 lifestyle and 14 health-related (see **Methods**)–using two complementary multivariable approaches. PERMANOVA evaluated differences in weighted UniFrac distances to test the overall effect of each variable, while EnvFit assessed contributions of these factors along the top three principal coordinate axes, helping to identify which factors most strongly contribute to community variation. We first applied a full model to 312 individuals with complete data for all 27 variables, identifying significant factors contributing to oral microbiome after adjusting for demography and lifestyle. To test robustness, we then applied a reduced model–retaining only the significant variables from the full model to a larger cohort of 618 individuals with available data. Across all these analyses, BMI consistently emerged as a significant factor associated with oral microbiome variation, highlighting it as a key variable for further investigation. Due to its high incidence in the UAE and its recognized clinical implications in the development of other cardiometabolic conditions^27^, we further investigated obesity’s contributions to the oral microbiome using multi-omics.

### Matched analysis highlights biomarkers of metabolic dysregulation in obesity

To mitigate potential lifestyle related confounding factors, we identified 97 obese individuals (BMI≥30) and 95 individuals within a healthy BMI range (controls,18.5≤BMI≤24.9) matched for demography, lifestyle, and oral health variables (**Methods**).

Principal Component Analysis (PCA) of clinical biomarkers, followed by multiple regression modeling (EnvFit) revealed that among all demographic and lifestyle variables, only obesity was significantly associated with variation in the PCA space (P=2.7×10^−3^, *EnvFit*, **Fig 2a,b**). This confirms that matching effectively minimized potential confounders. A random forest classifier further distinguished obese from control participants with moderate accuracy (OOB= 34%; AUC= 0.84), with with high variable importance scores (VIF) for several obesity-related serum markers that were elevated in obese individuals–including triglycerides, gamma-glutamyl transferase (GGT), alanine aminotransferase (ALT), and alkaline phosphatase (ALP), and blood glucose levels (GLUH), and HbA1c (**Fig. 2c.d**). Triglycerides, frequently elevated in obesity due to insulin resistance and increased hepatic lipogenesis, are a hallmark of metabolic syndrome^28^. Elevated liver enzymes (GGT, ALT, and ALP) reflect hepatic stress in obesity^29–31^, while increased glucose and HbA1c indicate impaired glycemic control and heightened risk for type 2 diabetes^32^. Elevated levels of key cardiometabolic biomarkers in the serum of the obese group emphasize the role of obesity as a major contributor to cardiometabolic risk.

**Figure 2.**
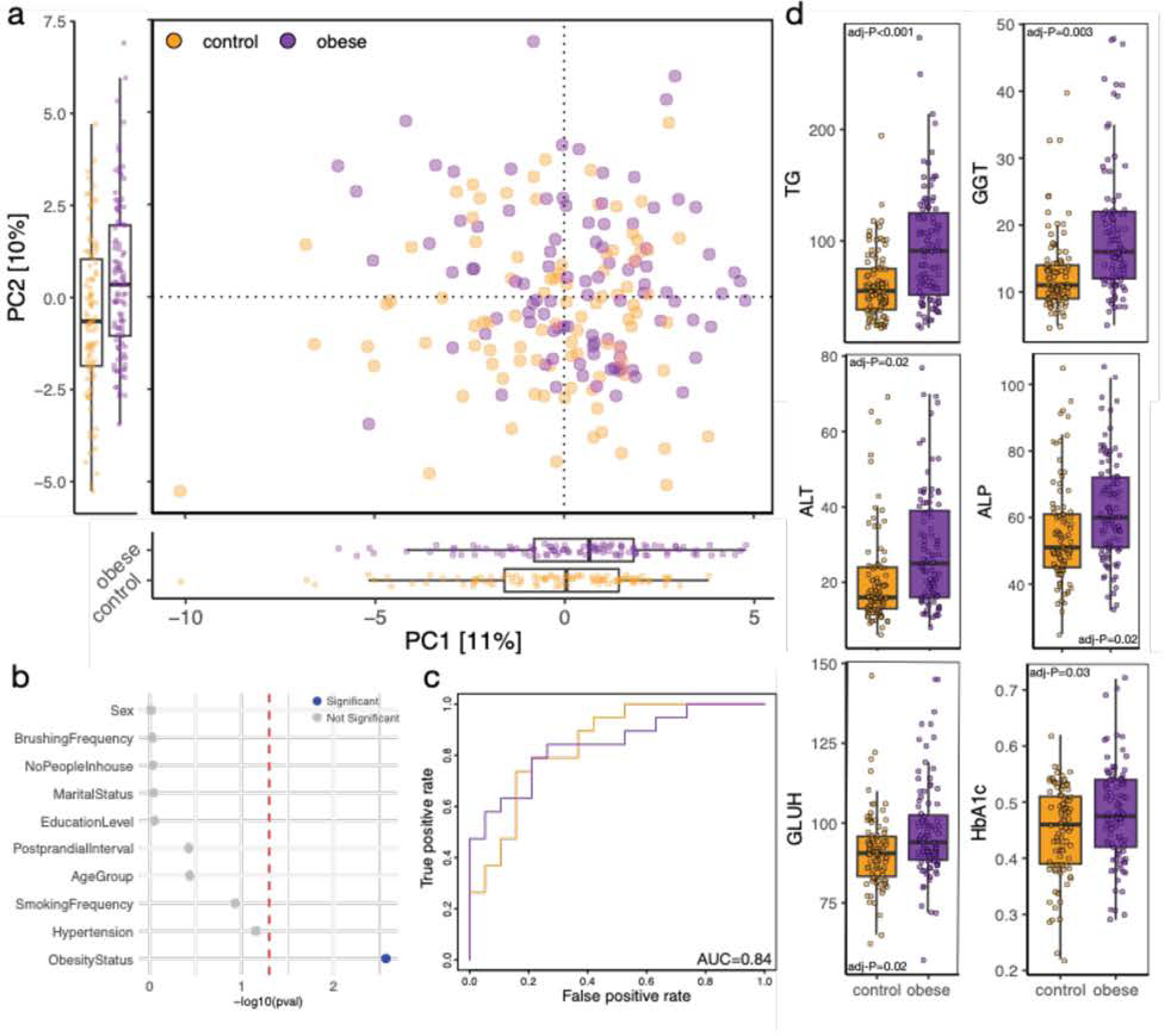
Obesity associated cardiometabolic biomarkers in the matched cohort. **a,** PCA of clinical markers revealed significant differences between the obese and controls along the top two principal components. **b,** p-values of demographic and lifestyle variables independent of the clinical markers. 13. **c,** AUC curves for random forest classifier of obese (purple) and control (orange) individuals; **d,** clinical markers with high VIF in the random forest analysis differ significantly between the obese and control groups. TG: triglycerides, GGT: gamma-glutamyl transferase, ALT: alanine aminotransferase, ALP: alkaline phosphatase, GLUH: blood glucose level, and HbA1c: Hemoglobin A1c.

### Obesity associated alterations in oral microbiome composition

To characterize oral bacteria at the species level and compare functional potential of the oral microbiomes between obese and control groups, we performed shotgun metagenomics sequencing of the 192 samples, generating ∼6.1 billion reads. After removing one sample with low sequencing depth, a total of 191 samples (obese n=97, controls n=94) remained. Each sample was sequenced with an average of 31.7 million reads, of which ∼3.7 million reads mapped to oral bacteria. Analysis of the microbial community structure, inferred by mapping individual reads to species-specific marker genes using MetaPhlAn4^33^, followed by Principal Coordinates Analysis (PCoA), revealed differences in oral bacterial composition between obese participants and controls (**Fig. 3 a**). A multivariate linear model implemented in MaAsLiN2^34^ revealed a total of 26 out of 366 bacterial species that were differentially abundant between obese and control participants (FDR adjusted P<0.05, **Fig. 3b**). A random forest model using these differentially abundant species effectively differentiated between obese and controls with appreciable accuracy (**Fig. 3c**, OOB=33%, AUC=0.75). Bacteria enriched in obesity included inflammation associated species such as *Streptococcus parasanguinis, S. gordonii,* and *S. infantis,* and *S. cristatus*^35^; as well as *Actinomyces oris,* and *A. massiliensis*, which are linked to a spectrum of chronic and infectious diseases^36^. The elevated presence of these taxa may create a microenvironment conducive to the growth of cariogenic bacteria, including *Lactobacillus gasseri* and *Limosilactobacillus fermentum*^37^; which— although less frequently detected–were more abundant in the obese participants.

**Figure 3.**
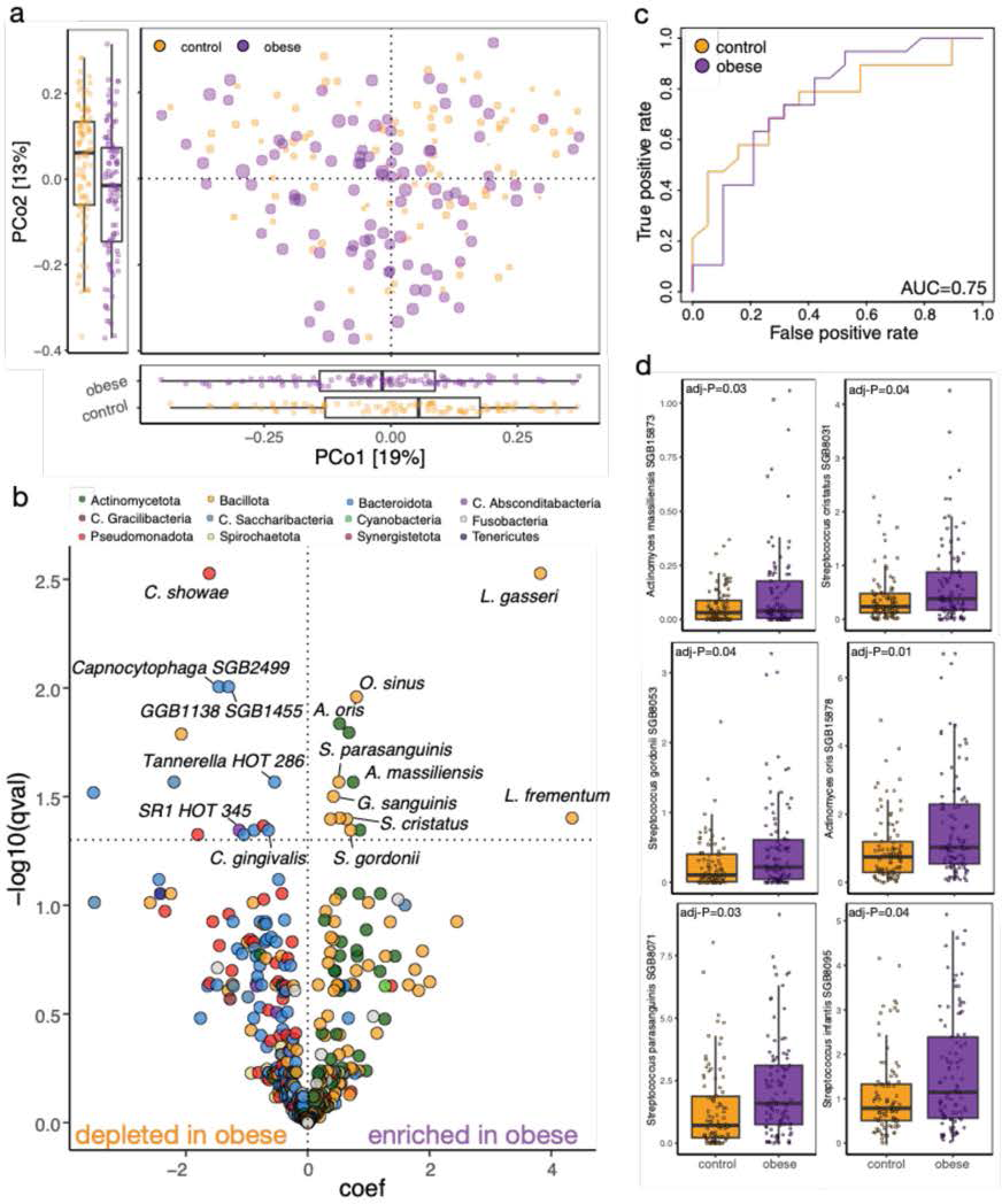
Obesity associated variation in the oral microbiota assessed using shotgun metagenomic sequencing. **a,** PCoA performed using Bray-Curtis distances at the species level for the 191 individuals (obese n=97, controls n=94) reveals small but significant differences in the oral microbiome composition between the obese and control groups. Each dot is an individual and color denotes obesity status (purple:obese, orange: controls). Size of the dot is scaled to the BMI of the individual. Boxplots reveal differences in the distributions of obese and control individuals across the two primary PCo axes. Both PCo 1 and 2 were associated with obese status (*P* = 0.035 and 0.0016 respectively, *GLM*). **b,** Volcano plot showing 26 differentially abundant species between obese and controls. Key species are labeled. **c,** Receiver Operating Characteristics (AUC) curve for a random forest classifier trained using the 26 differentially abundant species differentiates the obese (purple) from control (orange) individuals; **d,** Boxplots demonstrating differences in relative abundances of key obesity enriched species with FDR-adjusted P-values.

Notably, several species of bacteria that are typically found in the oral cavity such as *Capnocytophaga SGB2499*, *SR1 HOT 345 SGB6893*, and *Tannerella HOT 286 SGB2048* were depleted.

Amplicon sequencing confirmed these patterns. Specifically, obese participants showed a higher BBR (*P*=5×10^−4^, *GLM*), lower microbial diversity (*P*=0.031, *GLM*), and distinct microbial composition compared to controls (*P*<0.001, *EnvFit*). ASVs associated with with poor oral health, such as *Streptococcus, Actinomyces,* and *Lactobacillus* were enriched in obesity, while those linked to oral health, such as *Haemophilus, Capnocytophaga,* and *Veillonella*^18,38^, were depleted. These findings collectively underscore the distinct differences in various features of the oral microbial ecosystem in obesity, which are consistent with previous reports^17,18,20^.

### Obesity associated functional changes correspond to metabolic alterations

In addition to the shifts in oral microbial diversity and composition, we identified significant shifts in microbial metabolic capacity associated with obesity. Specifically, 94 out of 321 identified microbial metabolic pathways differed significantly between obese and control groups (**Fig 4a**, FDR-corrected *P*<0.05, *MaAsLin2*). At the enzyme level, 457 of the 1,824 microbial enzymes showed differential abundance between groups (FDR-corrected *P*<0.05) mirroring the observed pathway-level changes (**Figure 5**). Of the 94 obesity-associated pathways, 53 pathways were enriched in the obese group with 45 mapping to key metabolic functions–including fermentation of dietary carbohydrates into lactate, histidine metabolism into imodazole propionate (IMP), uracil/uridine biosynthesis, and the production of obesogenic metabolites–several of which have been previously implicated in obesity^39–47^ (**Fig. 5**). Plasma lactate is often elevated in obesity and is linked to inflammation, insulin resistance, and cardiometabolic risk^40–42^. Obesity associated microbial histidine degradation pathways lead to biosynthesis of urocanic acid and IMP, which impair insulin signaling^43,44^. Aminoacids such as cystine, methionine, and leucine are elevated in obesity and associated with adipogenesis in obesity^45–47^. In contrast, 41 pathways, including several involved in B-vitamin biosynthesis, were depleted in obesity.

**Figure 4.**
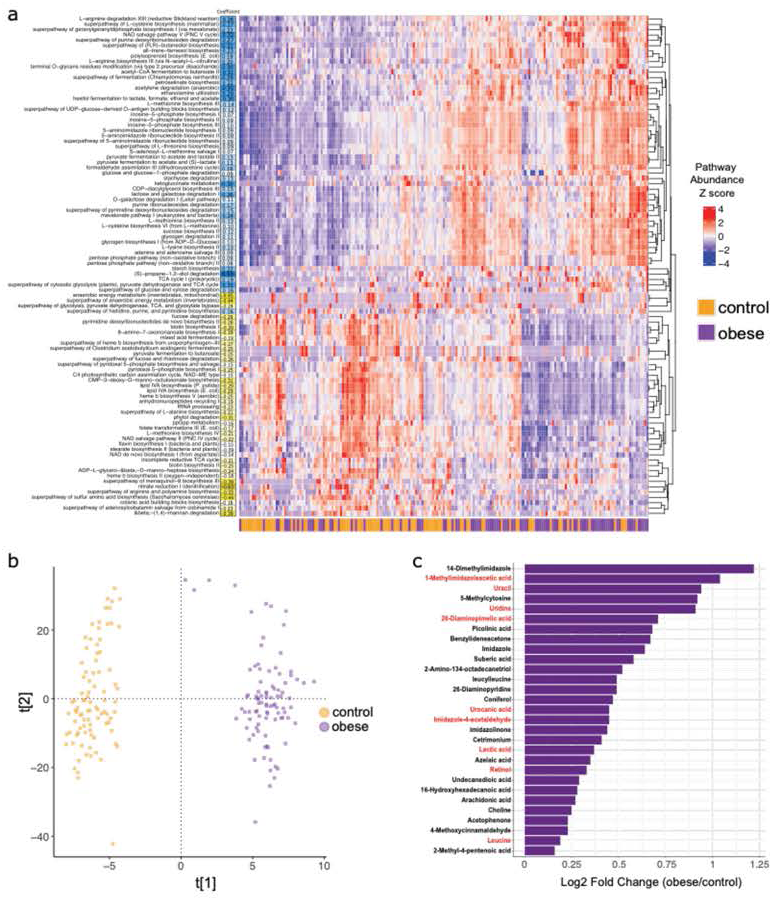
Obesity-associated oral microbial functional differences. **a,** Abundance of the 94 bacterial pathways significantly altered in obesity. The differentially abundant pathways are in rows, while each column represents an individual. The barplot below the heatmap indicates the obesity status of the individual. The numbers on the left column show the R^2^ values obtained from MaAsLiN2. Cells in blue indicate pathways enriched in obesity and those in yellow show obesity-depleted pathways. The dendrograms from unsupervised hierarchical clustering on the right cluster pathways according to obesity status. **b,** An orthogonal partial least squares-discriminant analysis (OPLS-DA) using the normalized abundances of all metabolites. Each circle is an individual, color coded by their obesity status. The orthogonal component and the top predictive component of the OPLS-DA are shown. The predictive component (t[1]) discriminates obese and control individuals. OPLS-DA cross-validation values were: R^2^X = 0.28, R^2^Y = 0.96 and Q^2^ = 0.38. **c,** Bar graph depicting the Log2 fold change of 29 metabolites enriched in obesity. Metabolites resulting from obesity enriched pathways and enzymes as shown in Fig. 5 are labeled in red.

**Figure 5.**
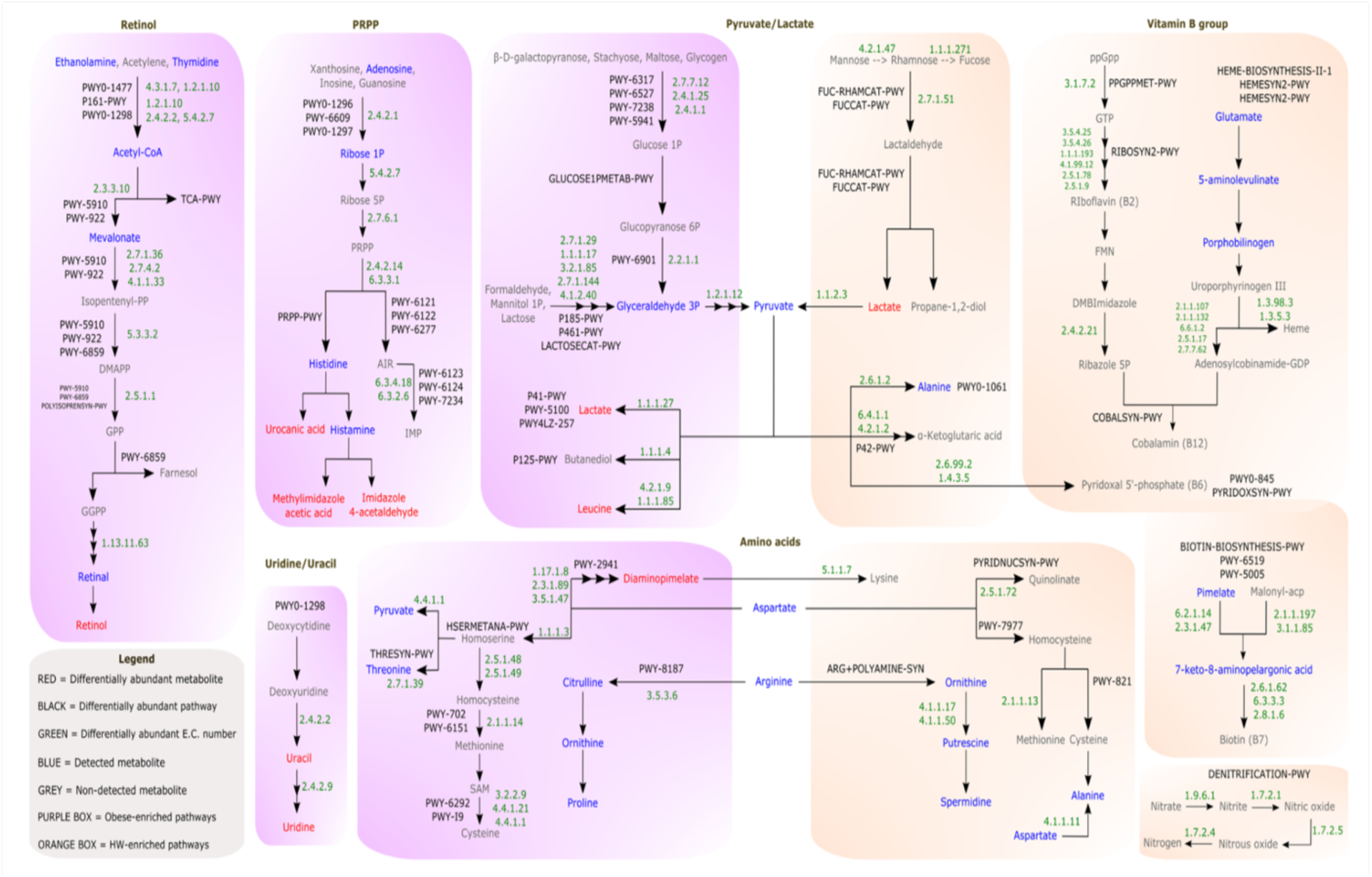
Obesity-associated oral microbial metabolic reprogramming. Of the 94 obesity associated pathways, 67 were involved in seven categories shown in boxes above. These microbial pathways (BioCyc; black text) and obesity-associated enzymes (E.C. numbers; green text) are shown. Pathways and enzymes in purple boxes are enriched in the obese group and those in orange boxes are depleted. Metabolites enriched in the obese are in red, their fold changes in obese relative to controls are shown in Fig. 4c. Blue indicates metabolites detected in the dataset but their abundances did not differ significantly between obese and control groups. Intermediate metabolites potentially involved in these pathways but not detected in our dataset are shown in gray. TCA= Tricarboxylic Acid Cycle; DMAPP=dimethylallyl pyrophosphate; GPP=geranyl diphosphate; GGPP=geranylgeranyl diphosphate; PRPP=5-phospho-α-D-ribose 1-diphosphate; AIR=5-aminoimidazole ribonucleotide; ppGpp=guanosine tetraphosphate; FMN=flavin mononucleotide; DMB=dimethylbenzimidazole; GMP=guanosine monophosphate; GTP=guanosine triphosphate; IMP=inosine monophosphate.

To investigate whether these microbial functional shifts contribute to metabolic changes in the oral environment, we performed untargeted LC-MS on all 191 samples and compared salivary metabolite profiles between obese participants and controls. To enrich for bacterial-derived metabolites, we centrifuged the saliva samples and discarded the supernatants, which typically contain host epithelial and dietary compounds. Metabolite profiling of the resulting bacterial cell pellets yielded 3,551 high-quality metabolite features after quality control (**Methods**). Multivariate analysis using orthogonal partial least squares-discriminant analysis (OPLS-DA) revealed clear separation between obese individuals and controls (**Fig 4b**).

Next, we annotated these features using curated bacterial metabolite databases, identifying 285 metabolites based on stringent criteria (**Methods**). These metabolites spanned various chemical classes, including organic acids, energy molecules, nucleotides, and carbohydrates. Importantly, they were strongly enriched for compounds commonly found in saliva and feces and showed no enrichment patterns consistent with dietary exposure. Notably, the two most significantly enriched pathways were histidine metabolism and arginine biosynthesis, both of which are canonical bacterial processes and not known to be active in oral epithelial cells^48^, further supporting their microbial origin.

To assess whether these metabolic shifts reflected underlying changes in microbial function, we performed a Procrustes analysis, comparing distance matrices of microbial functional profiles and metabolite profiles using the 285 annotated metabolites. This analysis revealed a high correlation (rho = 0.69, *P*=0.001, *permutation test*), suggesting that obesity-associated changes in the oral microbiome contribute to the observed metabolic differences. Consistent with this, 29 metabolites were found significantly different between the obese and control groups, all of which were enriched in the obese group (**Fig 4c**). Notably, nine of these metabolites correspond to known end products of microbial pathways and enzymes that were also enriched in the obese group (**Fig 5**), reinforcing the link between microbial metabolic activity and altered salivary metabolome in obesity.

The obese group showed enrichment of the superpathway of pyrimidine deoxyribonucleosides degradation (PWY0-1298), along with two key enzymes involved in uracil and uridine production: pyrimidine-nucleoside phosphorylase (E.C. 2.4.2.2) and uracil phosphoribosyltransferase (E.C. 2.4.2.9). Consistent with these findings, we also observed higher levels of salivary uracil and uridine in the obese group (**Fig 5**). *Lactobacilli*, which were also enriched in the obese group, can convert uracil and 5-phospho-α-D-ribose 1-diphosphate (PRPP) into uridine 5-monophosphate (UMP)^49^. UMP increases circulating uridine, which enhances hunger and higher caloric intake in humans^39^. Obese individuals also showed enrichment in pathways that convert nucleosides to PRPP. Key enzymes, including ribose-phosphate diphosphokinase (E.C. 2.7.6.1), amidophosphoribosyltransferase (E.C. 2.4.2.14), and 5-(carboxyamino)imidazole ribonucleotide synthase (E.C. 6.3.4.18), facilitate the conversion of PRPP into inosine 5-monophosphate (IMP), which plays a significant role in obesity-related insulin resistance^50^ and may contribute to non-alcoholic fatty liver disease^51^. Additionally, PRPP serves as a precursor in histidine biosynthesis. Notably, histidine is inversely associated with BMI^52^ and improves glycemic control when administered orally^53^, whereas its derivatives, 1-methylimidazole acetic acid^54^ and urocanic acid^55^—both of which are enriched in the saliva of our obese participants–are associated with obesity and obesity-related comorbidities (**Fig 5**).

Furthermore, the oral microbiome in the obese group was enriched for pathways related to lactate production, methionine metabolism, and farnesol biosynthesis (**Fig 5**). Pyruvate derived from metabolism of dietary sugars is catalyzed into lactate by L-lactase dehydrogenase (E.C. 1.1.1.27), an enzyme that was enriched in the obese group and correlated with higher levels of lactate in their saliva. Lactate accumulation in adipocytes is a key mediator of obesity-induced inflammation and systemic insulin resistance in mice^56^. Recent studies have linked elevated serum lactate levels with metabolic disorders^57^ and highlighted the potential role of adipose lactate as a signaling molecule in obesity^40^. Moreover, methionine metabolism has been associated with adiposity and weight gain^58^, while farnesol metabolism has been linked to sleep deprivation in obesity^59^. These findings collectively underscore the intricate connections between oral microbial pathways and metabolic dysregulation in obesity.

In addition, we detected an enrichment of retinol in the obese group, consistent with the enrichment of several pathways and enzymes related to the production of retinal, its precursor. Retinal is converted into retinol by beta-carotene 15,15’-dioxygenase (E.C. 1.13.11.63), which was also enriched in the obese participants (**Fig 5**). While the role of retinol in metabolic disease is emerging^60^, further studies are needed to clarify its role in obesity. Similarly, we detected elevated levels of diaminopimelate (DAP) in the obese group, which results from catalysis of aspartate by several enzymes (e.g. E.C. 1.17.1.8; 2.3.1.89; and 3.5.1.47) within the L-lysine biosynthesis pathway (PWY-2941). In contrast, controls were enriched for diaminopimelate epimerase (E.C. 5.1.1.7), which converts DAP into lysine. The direct association between DAP and lysine with obesity is not well understood, although lysine has been shown to reduce body weight in mice fed a high-fat diet^61^. The roles of these molecules in obesity warrant further investigation.

Obese participants also showed a depletion of 41 pathways, many of which are involved in the biosynthesis of essential B-vitamins, including flavin (B2), pyridoxal 5-phosphate (B6), biotin (B7), and cobalamin (B12), as well as pathways involved in heme synthesis and nitrate reduction (**Fig 5**). B-vitamins are crucial for numerous human physiological processes and the proper functioning of microbial ecosystems^62^. Notably, considering precursors for Vitamin B12 biosynthesis, enzymes involved in the conversion of guanosine triphosphate to riboflavin (e.g. riboflavin synthase; E.C. 2.5.1.9), the subsequent conversion of dimethylbenzimidazole to α-ribazole-5’-phosphate (e.g. nicotinate nucleotide dimethyl benzimidazole phosphoribosyl transferase; E.C. 2.4.2.21), and production of adeonsylcobinamiede-GDP from uroporphyrinogen (e.g. uroporphyrinogen-III C-methyltransferase; E.C. 2.1.1.107 and adenosylcobinamide-phosphate guanylyltransferase; E.C. 2.7.7.62), were depleted in obese participants. Vitamin B12 and heme biosynthesis pathways share several enzymes^63^, and two enzymes involved in converting uroporphyrinogen to heme (e.g. protoporphyrinogen IX dehydrogenase; E.C. 1.3.5.3 and coproporphyrinogen dehydrogenase; E.C. 1.3.98.3), were also depleted in obese participants. B12 deficiency can alter host lipid metabolism and lower heme levels are linked to adipogenesis^64^, both contributing to obesity^65^. Additionally, vitamins B6 and B7 play key roles in energy production via carbohydrate and lipid metabolism^66^, potentially mitigating the severity of obesity^67^. Since humans cannot synthesize most vitamins, they rely on gut bacteria to generate these essential nutrients from dietary sources^62^. Our results suggest that the depletion of B vitamin producing bacteria from the oral cavity could impact energy metabolism, contributing to obesity, obesity-related comorbidities, and poor overall health^67,68^. To our knowledge, no publicly available oral microbiome datasets currently include both metagenomic and metabolomic data, limiting opportunities for external replication. Nonetheless, these findings lay a strong foundation for future functional validation via strain isolation and experimental assays.

### The oral bacterial species contributing to obesity associated functional differences

To evaluate the contribution of key species to microbial functions in obesity, we assessed potential correlations between differentially abundant bacterial pathways and the top 20% most abundant bacterial species. While no single bacterial species was exclusively associated with the 94 obesity-associated pathways, 7 out of the 26 bacterial species enriched in obesity correlated positively with obesity-enriched pathways and exhibited a strong negative correlation with the pathways depleted in obesity (FDR adjusted p<0.05, *Spearman’s Rank Correlation Test*). For instance, several pathways related to lactate production and purine or pyrimidine nucleoside metabolism were positively correlated with obesity enriched bacteria, including *Gemella sanguinis*, *A. oris*, *S. parasanguinis*, *S. cristatus*, and *S. gordonii* in the obese group. Conversely, these obese-enriched bacterial species showed a strong negative correlation with pathways related to B-vitamins.

Next, we generated 1,576 metagenome-assembled genomes (MAGs) from 191 samples, which included 810 MAGs spanning 179 taxa from the obese individuals and 766 MAGs spanning 192 taxa from the controls. After dereplication, we retained 221 unique MAGs (uniMAGs, ANI≥95%) of medium or higher quality (based on MIMAG standards^69^, >50% completeness and <10% contamination on CheckM^70^). Forty-three of the 221 uniMAGs were not assigned to known species using the GTDB-tk v2 database (GTDB Release 226)^71^ and lacked a representative in UHGG ver2^72^. These MAGs, belonging to diverse phyla, potentially represent novel bacterial species (novMAGs) (**Fig. 6a**). Most novMAGs belonged to the phyla Bacillota, Pseudomonadota, and Patescibacteria. Their global distribution revealed monophyletic distant branches in relation to the reference genomes from their corresponding groups, suggesting differences in gene content compared to the representative genomes (**Fig. 6b**).

**Figure 6.**
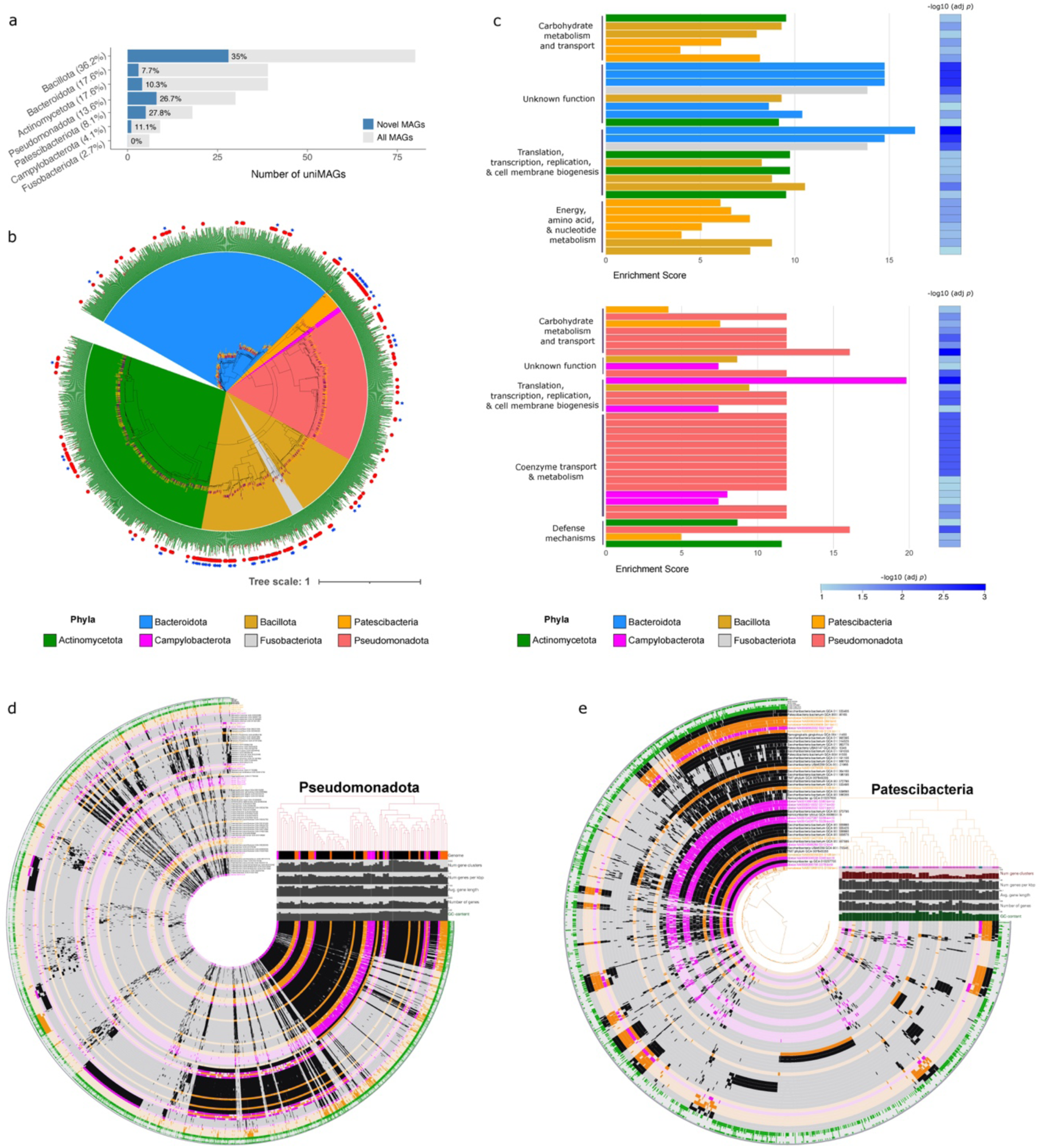
Taxonomic and functional diversity of MAGs recovered from obese and control individuals. **a,** A total of 221 uniMAGs of medium quality or higher (> 50% completion, <10% contamination) and according to MIMAG standards were recovered. The bars indicate uniMAGs obtained from each phylum and their percentages per phyla is shown in parenthesis. Blue bars denote novel MAGs and their percentages are listed to the right. Bacillota, Bacteroidota, and Actinomycetota had the most uniMAGs while Bacillota had the highest percentage of novMAGs (35%, n=28). **b,** The phylogenomic relationship between the 1,576 MAGs based on 74 single copy gene-set Hidden Markov Models (HMMs). The outer blue stars indicate novel MAGs and the red dots indicate uniMAGs. The outer bars indicate completeness (green) and contamination (red) for each MAG. Phyla are colored according to the legend. Branch colors indicate MAGs obtained from obese (purple) and control (orange) participants. **c,** COG and KEGG functions enriched in the MAGs obtained from the obese (top) and control (bottom) groups. In the obese group, a notable number of enriched genes with unknown functions came from Bacteroidota MAGs and MAGs from Bacillota were enriched for genes involved in energy metabolism. Whereas MAGs from controls showed enrichment of coenzyme transport and metabolism category, which included genes involved in cobalamin (vitamin B12) biosynthesis. The categories Translation, Ribosomal structure and biogenesis, Transcription, Replication, Recombination and repair and cell wall/membrane/envelope biogenesis were grouped together into one category called Translation, transcription, replication, & cell membrane biogenesis. **d-e,** Pangenomic analysis for Pseudomonadota **(d)** and Patescibacteria **(e)** including dereplicated MAGs from obese (purple), controls (orange), and reference genomes (black). Gene clusters are organized according to a hierarchical clustering of their presence or absence (central dendrogram). Genomes are organized based on the presence or absence of gene clusters (top right dendrogram). Opacity denotes presence or absence of the COG function in each genome.

Pangenome analysis of the uniMAGs along with their closest taxonomic representatives for gene clusters obtained from Clusters of Orthologous Groups (COG) and the Kyoto Encyclopedia of Genes and Genomes (KEGG) databases^73,74^ further revealed significant enrichment of COG functions and KEGG modules related to coenzyme transport and metabolism in MAGs from controls (FDR-corrected *P*<0.05, **Fig. 6c**). Specifically, several genes involved in cobalamin (vitamin B12) biosynthesis were enriched in Pseudomonadota MAGs obtained from the controls (FDR-corrected *P*<0.05, **Fig. 6c**,**d**) while Patescibacteria MAGs obtained from obese participants were enriched for genes involved in histidine degradation and uridine monophosphate biosynthesis (FDR-corrected *P*<0.05, **Fig. 6c**,**e**).

Previous metagenomics studies have detected cobalamine synthesis genes in diverse bacteria, including species common in gut^75^ and oral comunities^63^, which is consistent with our results. These results suggest that altered functional potential of the oral microbiota may contribute to metabolic perturbations in obesity.

### Integrative multi-omic analysis reveals microbe-metabolomic signatures linked to obesity-associated clinical markers

Associations between shifts in the microbiome, metabolites, and clinical phenotypes indicate that changes in the oral microbiome may significantly impact host physiology and vice versa. A comprehensive association analysis integrating microbiome, metabolite, and serum cardiometabolic biomarkers revealed plausible biological mechanisms by which oral bacteria may contribute to obesity. Bacteria-metabolite correlations revealed obesity-enriched species such as *O. sinus* were positively associated with acidic metabolites including lactate, which plays a major role as a signaling molecule in obesity^40^ leading to metabolic disorders^57^. Accordingly, lactate and these microbial species were positively correlated with serum triglycerides **(Fig. 7a)**. Acidic metabolites may lower oral pH, which favors lactic acid bacteria growth. *Streptococcus* species, including *S. parasanguinis*, harbor a glycerophosphodiesterase enzyme (glpQ) that can metabolize dietary glycerophosphocholine into choline and glycerol-3-phosphate^76^. Choline is essential for survival and is incorporated into the bacterial cell wall by biofilm producing bacteria in the oral cavity^77^, while glycerol-3-phosphate is a triglyceride precursor. Consistent with these functions, *S. parasanguinis* and several other obesity-enriched bacterial species were positively associated with glycerophosphocholine, choline, and serum triglycerides (**Fig. 7a,b**).

**Figure 7.**
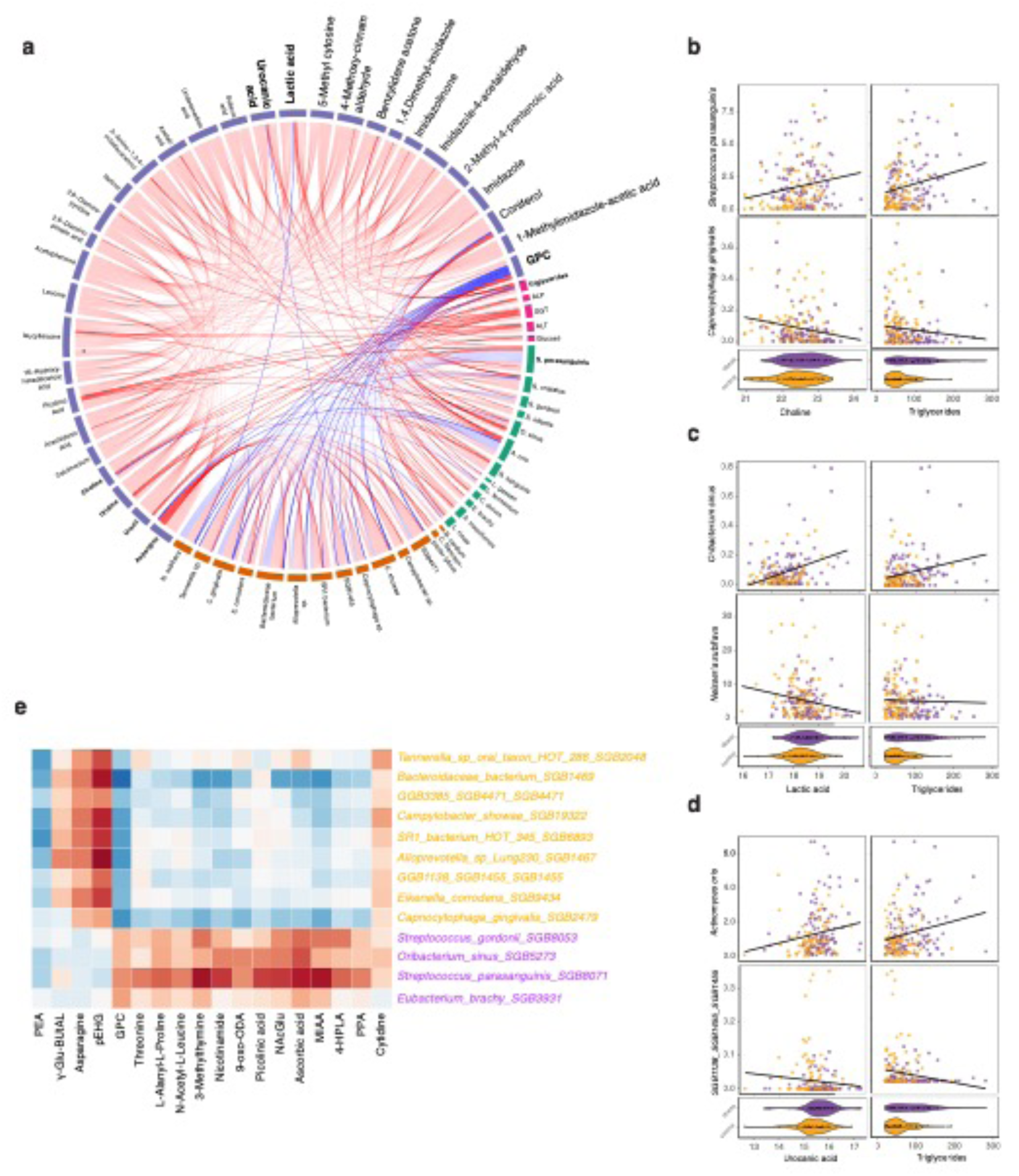
Microbiome-metabolite shifts are associated with serum cardiometabolic markers. **a,** Correlations between obesity-associated microbial species, metabolites, and cardiometabolic serum markers. Red lines indicate significant positive correlations, and blue lines represent negative correlations. Several obesity-enriched bacterial species (green outer segments) are positively associated with obesity enriched metabolites (blue outer segments) as well as serum cardiometabolic markers (pink outer segments). Obese depleted species (brown outer segments) are negatively associated with these metabolites and serum markers. **b,** Scatter plots showing positive associations between *S. parasanguinis*, choline, and serum triglycerides (top row), while *C. gingivalis* is negatively associated with both (bottom). **c,** *O. sinus* is positively associated with lactate and triglycerides (top). *N. subflava*, a common oral bacterium, has negative associations with both of these features (bottom). **d,** *A. oris,* urocanic acid, and triglycerides show positive correlations, while *SGB1455*, an uncharacterized species which is depleted in the obese is negatively associated with both (bottom). Violin plots demonstrate differential abundances of these metabolites and triglycerides between obese and controls. **e,** Key bacteria-metabolite associations identified by HAllA. PEP: Phosphoethanolamine, γ-Glu-ButAL: N-(4-oxobutyl)-L-glutamine, pEHG: pyroglutamyl glycine, GPC: glycerophosphocholine, 9-oxo-ODA: 9-oxo-10(e) 12(e)-octadecadienoic acid, NAcGlu: N-Acetyl-L-glutamic acid, MIAA: Methylimidazoleacetic acid, 4-HPLA: 4-hydroxyphenyllactic acid, PPA: phenylpyruvic acid.

Moreover, lactic acid bacteria possess histidine decarboxylase enzymes capable of breaking down histidine into imidazoles^78^. Consistent with this, *S. parasangiunis*, *A. oris*, *L. gasseri*, and *L. fermentum* were also positively associated with histidine and pyrimidine degradation pathways as well as their metabolic products such as imidazoles, urocanic acid, uracil, and uridine (**Fig. 7a,d**). Uracil and uridine regulate liver functions^79^ and urocanic acid, along with other histidine-derived metabolites, are linked to obesity related comorbidities^54,55^. Consequently, both uridine and uracil were associated with elevated ALP levels and urocanic acid correlated with elevated triglycerides. Conversely, common human oral cavity residents and obesity-depleted bacterial species such as *Neisseria subflava*, *C. gingivalis*, and *SGB1455* showed an inverse correlation with obesity-associated metabolites as well as molecular biomarkers (**Fig. 6b-e**). Instead, several of such bacteria were positively correlated with asparagine, a metabolite inversely associated with obesity in Middle-Eastern adults^80^.

We also evaluated bacteria-metabolite associations using hierarchical-all-against-all association testing (HAllA)^81^. This machine learning framework identifies covarying species-metabolite pairs in high-dimensional datasets and is well suited for detecting non-linear associations and handling sparsity, making it more robust than traditional correlation based methods. HAllA corroborated positive associations between *O. sinus* and *Streptococcus* species and acidic metabolites (**Fig. 7e**). In addition to corroborating microbiome-metabolite pairs described above, HAllA identified several additional bacteria-metabolite associations that are relevant for oral microbial ecology (**Fig. 7e)**. For example, we found strong positive associations between several obesity-depleted bacteria and pyroglutamyl glycine, which serves as a nutrient source or signaling molecule for bacteria, can modulate protein function and stability through post-translational modifications^82^. Phosphoethanolamine was negatively associated with both obesity-enriched and depleted oral taxa, likely reflecting its broad utilization by oral bacteria for membrane biosynthesis and cell envelope remodeling.

Collectively, these results indicate that obesity-enriched oral bacteria change the oral environment and generate metabolites that can interfere with distant organ functions. In addition to these, we detected many more bacteria-metabolite-serum metabolic marker associations in this dataset, which may be relevant to obesity and warrant further exploration.

## Discussion

This study represents the first and most comprehensive analysis of oral microbial perturbations in obesity to date, addressing critical gaps in understanding the role of the oral microbiome in systemic health. Previous studies have been limited by small sample sizes, inconsistent microbiome profiling and analysis techniques, and inadequate taxonomic resolution, making it challenging to interpret their results. For example, studies comparing obese and normal weight individuals in the US (14 obese, 19 controls) and Qatar (37 obese, 36 lean) showed no significant differences in oral microbial diversity and composition, likely due to insufficient sample sizes^83,84^. Although a larger study in 647 obese and 969 non-obese Americans corroborated these results^38^, a Chinese cohort (n=659) identified significant associations between BMI and beta diversity^85^. These inconsistencies, partly due to differences in microbiome profiling and bioinformatics techniques, underscore the need for large-scale, high-resolution studies. Additionally, lifestyle factors such as diet may contribute to discrepancies in findings, although such differences tend to be small compared to the gut microbiomes^86^. Furthermore, because of the use of 16S rRNA, oral microbiome analysis of most of these studies is limited to broad taxonomic levels. At the phylum level, Bacillota-to-Bacteroidota ratio is positively associated with obesity^84,87^ along with higher relative abundances of *Streptococcus*, *Gamella*, *Oribacterium*, *Corynebacterium*, *Bifidobacterium*, *Lactobacillus*, and *Actinomyces*^38,83^.

In this study, we generated a large, well-characterized, and deeply phenotyped cohort to overcome these challenges and provide new insights into the oral microbiome’s role in obesity. We observed a higher Bacillota-to-Bacteroidota ratio in obese participants, along with significant differences in microbial diversity and composition. Notably, pro-inflammatory species such as *Streptococcus parasanguinis, S. gordonii, S. cristatus*, *Gamella sanguinis*, *Oribacterium sinus*, *Corynebacterium durum*, *L. gasseri*, and *Actinomyces oris* were enriched in obesity, while several novel species of bacteria that are typically found in the oral cavity such as *SR1 HOT 345 SGB6893*, *Capnocytophaga SGB2499, Tannerella HOT 286 SGB2048, Bacteroidaceae bacterium SGB1469*, *GGB3385 SGB4471*, and *GGB1138 SGB1455* were depleted. Many of these microbes were also associated with serum metabolic markers diagnostic of obesity-related metabolic reprogramming, including elevated triglycerides and ALP, indicaing systemic inflammation in obesity.

All three obesity enriched *Streptococcus* species are natural members of the oral flora, playing a key role in oral biofilm formation and shaping the oral microbial ecosystem through quorum sensing. Their metabolic activities include fermenting dietary carbohydrate, which produces lactic acid and acidifies the oral cavity, creating an environment where opportunistic bacteria like *S. parasanguinis* can thrive. Chronic acidification weakens the oral mucosal barrier by degrading mucin and disrupting the integrity of the epithelial layer, allowing metabolites produced in the oral cavity to enter the blood and reach distant organs^13,88^. Consistent with this, we found an enrichment of *Oribacterium sinus*, *L. fermentum, and L. gasseri* in the obese participants. These bacteria are known to produce lactic acid as an end product^89^. Obese participants were characterized by elevated levels of lactic acid, which upon entering the bloodstream, can increase triglyceride levels and trigger adipogenesis^40,56^, which are key features of obesity^57,90^.

Although some *Streptococci* are part of the normal oral flora, *Streptococcus parasanguinis, S. gordonii,* and *S. cristatus* have been linked to systemic inflammation in obesity and metabolic syndromes^35^. These bacteria contribute to the metabolism of glycerophosphocholine into choline and triglycerides, both of which are key factors in obesity-related metabolic disturbances and poor cardiovascular health outcomes^91^. Notably, the Emirati diet, rich in animal products such as eggs, meat, fish, and dairy, provides phosphatidylcholine and histidine, which are converted by microbial enzymes in the into metabolites such as TMA, TMAO, glycerophosphocholine, and choline^92^. Although these processes are usually attributed to the gut microbes, our results revealed that several of these obesity enriched bacteria are positively correlated with glycerophosphocholine, choline, and elevated serum triglycerides. This suggests that oral bacteria play a critical role in regulating host metabolism through diet, ultimately influencing cardiometabolic health.

Beyond lactic acid production, obesity-enriched bacteria, such as *L. fermentum*, are major histidine degraders and produce histamine, which is further converted into urocanic acid and imidazole compounds. Histidine is a crucial metabolite involved in immune regulation, protein synthesis, energy production, cellular metabolism, and overall homeostasis in the human body^93,94^. However, the degradation of histidine into histamine and imidazole compounds has been linked to inflammation, impaired appetite regulation, circadian rhythm disruption, and impaired insulin signaling^91^. Although histidine metabolism is commonly attributed to gut microbes, we observed several microbial pathways and enzymes involved in histidine metabolism were elevated in the oral microbiomes of the obese participants. This corresponded with higher levels of imidazole metabolites, indicating an increased histamine turnover, which could reflect an inflammatory response or metabolic dysregulation that is often seen in obesity^95^.

Furthermore, we also detected several metabolic pathways enriched in the obese group that link dietary carbohydrate metabolism with energy regulation. For example, carbohydrates from the diet can be converted into ribose-5-phosphate, which is then transformed into PRPP, which serves as a precursor for uracil and its derivative, uridine. Several pathways contributing to PRPP synthesis–including PRPP synthetase (E.C. 2.7.6.1), the key enzyme catalyzing PRPP formation–were elevated in the obese group.

Additionally, the superpathway of pyrimidine deoxyribonucleosides degradation (PWY0-1298), which potentially produces uracil and uridine, was also elevated in obese participants. Both uracil and uridine play a critical role in energy homeostasis^79^, promoting increased hunger and higher caloric intake^39^. Elevated levels of these metabolites have been associated with hyperglycemia and the generation of reactive oxygen species in obese rats^96,97^. Although the role of gut microbiome in energy metabolism is well established^98^, our findings suggest that disruptions in the oral microbiome may also contribute significantly to imbalances in energy homeostasis. These disruptions may trigger inflammatory responses and contribute to obesity-related comorbidities^93–95^.

In addition, several microbial pathways involved in B-vitamin synthesis were depleted in obesity, which may further contribute to metabolic dysfunction^99^. Previous studies have shown an inverse relationship between vitamin B12 and obesity^100^, while others have highlighted the role of gut microbiome in vitamin synthesis^101^. However, previous metagenomic studies have detected cobalamin biosynthesis genes and salvage pathways in a wide range of bacteria^63^. Yet, the link between the oral microbiome and vitamin B12 biosynthesis remains largely unexplored. We observed that Pseudomonadota MAGs derived from the controls have vitamin B12 biosynthesis capacities. In particular, we find an enrichment of B vitamin related KEGG functions in control derived *Lautropia mirabilis*, *Lautropia dentalis*, and *Ottowia sp*., which are commonly found in the oral cavity. Moreover, the adenosylcobalamin salvage pathway was depleted in the obese group, potentially reducing the bioavailability of vitamin B12. While this warrants further investigation, our results suggest that a small fraction of oral bacteria may contribute to local production of vitamin B12 in the mouth. Depletion of these bacteria in obesity may lead to impaired heme synthesis, which is essential for oxygen sensing and electron transfer^67^ in red blood cells^66^. A reduction in heme may lead to accumulation of fatty acids and exacerbate obesity^68^.

While previous studies have extensively linked the gut microbiome to obesity^6,102–104^, limited research has explored the contribution of the oral microbiome to systemic diseases. This gap provided an opportunity to leverage the mouthwash samples, collected as part of the UAEHFS, a result of years of efforts in overcoming cultural barriers through extensive community engagement. Paired with blood, urine samples, and detailed lifestyle information, these mouthwash samples allowed us to integrate high-dimensional multi-omic datasets to investigate oral bacteria, their metabolic activities, and their interactions with cardiometabolic markers. We focused on obesity due to its global relevance as a significant risk factor for cardiovascular diseases^1,2^ and its high prevalence among young Emirati nationals^5^.

Despite our best efforts, this study has several limitations that must be noted. First, we were unable to collect reliable dietary data from participants, highlighting the need for increased awareness regarding the role of diet in chronic diseases. We were also unable to completely eliminate the effect of smoking and post-prandial intervals, which affect both the microbiome and metabolome. Nevertheless, we identified numerous bacterial species, pathways, and specific enzymes that differed between obese and controls. These differences correlated with variations in respective metabolites as well as clinical parameters in the predicted directions, supporting microbiome-metabolite links in obesity. Second, a large portion of our sequencing reads mapped to the human genome, which likely hinders large-scale oral metagenomics research. Only 10% of the metagenomics reads were bacterial, limiting our ability to assemble MAGs representative of the oral microbiome. Despite these challenges, we assembled ∼1500 metagenomes from ∼200 bacterial species. Although this is fewer than expected based on 16S rRNA profiling from the same samples, we identified 43 MAGs (<0.5% of all MAGs) that had no close representative taxa in public databases, highlighting a substantial gap in our understanding of the taxonomical and functional diversity of human microbiomes, especially within underrepresented populations^105,106^. Finally, our metabolome likely represents only a subset of metabolites–primarily those most stable–due to sample collection limitations that may not have preserved volatile or less-stable metabolites. We also did not detect B vitamins in our metabolome, likely because saliva is a rich source of vitamin binding proteins secreted by salivary glands that bind free vitamins^107^. While these proteins may protect it from degradation in stomach, protein bound B vitamins in saliva samples is not detectable in the metabolome using LC-MS.

However, the oral metabolomic profiles we generated align closely with those from two previous studies involving small cohorts of patients with Type 2 diabetes mellitus (n=20)^70^ and depression-obesity comorbidity, though those studies lacked metagenomic data^71^. Future work focusing on improved sample preservation, depletion of human DNA, and deeper sequencing for better characterization of the oral microbiome will be essential to overcoming these limitations.

Our study utilizes an integrative framework to demonstrate alterations in oral microbiome composition, function, and metabolic activities of the oral microbiome in obesity. We offer evidence that obesity-related changes disrupt the carbohydrate and energy metabolic landscapes. Collectively, our findings expand current knowledge regarding the critical role of oral microbiome in obesity and offer initial insights into how bacterial cross-talk in the oral cavity may influence systemic disease beyond the oral health. Further research, especially studies incorporating diverse lifestyles, is needed to validate these findings on a global scale. Mechanistic investigations through longitudinal studies will offer a more comprehensive understanding of host-microbiome interactions and deepen our understanding into the biology of obesity.

These findings also hold potential for advancing microbiome-based prevention strategies and therapeutics to combat the global obesity epidemic. Finally, the robust associations between oral microbiome alterations and obesity in our cohort suggests the need to explore the role of oral microbiome in other complex diseases.

## Materials and Methods

### Ethics statement

The UAE Healthy Future Study (UAEHFS) was conducted according to the guidelines of the Declaration of Helsinki, and the study protocol was approved by the Institutional Review Board of Abu Dhabi Department of Health, reference number DOH/HQD/2020/516 and NYUAD’S Research Ethics Committee (REC, REF:0072017R). All participants read and understood the information leaflet and signed the consent form prior to their recruitment.

### Study design and mouthwash sample collection

The UAEHFS is a cross-sectional prospective cohort study designed to investigate the biological mechanisms underlying chronic diseases in Emirati nationals. Mouthwash samples were collected with written consent between 2016 and 2020, as described previously in detail^24^. Participants refrained from eating at least 1 hour prior to sample collection. Briefly, participants swished 10 mL of 0.9% sterile saline vigorously for 30 seconds and spit into sterile barcoded tubes. These tubes were immediately transported to the lab and stored at −80°C until analysis. We conducted a pilot study to compare the oral microbiome from saliva versus mouthwash samples collected using this protocol. Our results show that mouthwash is as reliable as saliva in capturing the oral microbiome, which is consistent with previous studies^108^.

In addition to providing mouthwash samples, participants underwent physical and clinical exams, which included anthropometric measurements, blood pressure assessments, blood and urine tests, and self-completed a demographic and lifestyle questionnaire. Clinical parameters were used to define prevalent risk conditions within the cohort based on established criteria. Participants were categorized based on body mass index (BMI) as underweight (BMI < 18.5), healthy weight (18.5 ≤ BMI ≤ 24.9), overweight (25 ≤ BMI ≤ 29.9), or obese (BMI ≥ 30) categories.

### Blood and urine biomarker measurements

Participants provided non-fasting blood and urine samples, which were analyzed for routine chemistry using the Beckman DXC600 (Beckman Coulter, USA) as described previously^24^. This provides us with biomarkers indicative of lipids, glycemic levels, liver and kidney function, serum electrolytes, and red blood cell indices.

### DNA extraction

The mouthwash samples underwent mechanical lysis using a Bead Ruptor 96 (Omni International, USA) followed by DNA extraction using the QIAGEN DNeasy Powersoil Pro Kit. To minimize cross-contamination, DNA extraction was performed in individual tubes, with a negative control, containing only the extraction reagents but no mouthwash, included after every 4-5 samples. This process yielded 669 mouthwash samples and 134 controls. DNA was eluted in nuclease-free water, and the concentration was measured using NanoDrop™ 2000 spectrophotometer (Thermo Scientific, USA). The DNA extraction protocol used in this study is available at DOI: https://dx.doi.org/10.17504/protocols.io.eq2ly7dyrlx9/v1.

### 16S rRNA gene sequencing

DNA amplification targeting the V4-V5 region of the 16S rRNA gene was performed on all samples and controls using the universal primer set 515F/926R^109^. Forward primers were barcode-tagged for sample multiplexing. PCR products were visualized on an agarose gel, purified using AMPure beads (Beckman, USA), and quantified with the Qubit BR kit (Thermo Scientific, USA). No PCR bands were observed in any of the controls. Thus, these were excluded from further steps. The PCR products from the samples were normalized, pooled, and sequenced on an Illumina MiSeq to generate 2×300 paired-end reads.

### 16S rRNA amplicon processing

A total of 8,850,022 paired-end reads were obtained from 669 samples. Quality assessment of the sequenced reads using FastQC revealed poor quality in the reverse reads, preventing their merge with the forward reads. Consequently, only forward reads were used for further analyses. Forward reads were processed with an in-house pipeline based on a previous study^110^, which implements DADA2^111^ to learn error rates, dereplicate amplicons, remove chimeras, identify amplicon sequence variants (ASVs), and construct a sequence table. The reads were trimmed to 150 bp to retain high quality sequences (Phred score >30). Reads containing N nucleotides or >2 expected errors were discarded (maxN = 0, maxEE = 2, truncQ = 2). ASVs were inferred from 5,936,465 high-quality reads, representing 67% of the initial dataset. Taxonomy was assigned using the RDP classifier^112^ utilizing the SILVA v138 training set as the reference. Multiple sequence alignment was conducted with DECIPHER^113^, and a maximum likelihood tree was built using the neighbor-joining method in phangorn^114^. Subsequent analyses were conducted in R v4.2.0 using the phyloseq package^115^. ASVs present in less than 5% of the samples were removed, leaving 5,851,506 reads that were assigned to 384 taxa. Three ASVs were identified as Eukaryota and one was unresolved at the phylum level. These four ASVs were excluded, resulting in 5,712,557 reads across 380 ASVs for subsequent analyses.

### Identification of matched cohort and lifestyle and health-associated variable selection

Out of the 669 participants in our study, we identified 97 individuals with obesity (BMI ≥ 30) and established a comparison group of 95 healthy weight controls (18.5 ≤ BMI ≤ 24.9). To ensure the comparability of these groups, we first conducted a linear regression analysis to identify variables significantly associated with BMI. We removed highly multicollinear variables from this regression (VIF> 10). We then used MatchIt v4.5.4 to match for sex, age, and other variables significantly associated with oral health. This ensured equal distribution of potential confounding variables that could influence the oral microbiome, such as postprandial interval, brushing frequency, and smoking frequency across both case and control groups.

### Blood and urine marker analysis

Principal component analysis (PCA) was performed on a set of 49 non-redundant blood and urine biomarkers, selected based on the availability of complete data across all 191 individuals in the matched cohort. PCA was conducted using the prcomp function in R. In an initial PCA, sex showed the strongest association with biomarker variation; therefore, we regressed out the effect of sex using the *limma* package and conducted all downstream analyses on the residualized biomarker data. Associations between metadata and biomarker composition were assessed using the *envfit* function. To evaluate the discriminatory power of biomarkers between control and obese individuals, a random forest classifier was trained on the residualized data, and performance was assessed using area under the receiver operating characteristic curve (AUC). Predictors with high variance inflation factors (VIF) were identified and retrieved from the model.

### Metagenomics library preparation and sequencing

We performed shotgun whole metagenomics sequencing for the 97 obese and 95 matched controls (n=192) at New York University Abu Dhabi Core Technology Platform Sequencing Center (Abu Dhabi, UAE) using Illumina DNA Library Preparation kit (Illumina, USA), following the manufacturer’s instructions using 300-450 ng of DNA in 30 µL volume. During the library preparation, a unique 10 base pair (bp) dual-indexed barcode was added to each sample. Sequencing libraries were size selected using AMPure XP beads (Beckman, USA) targeting a fragment length of 450 bp with insert size of 350 bp and checked using the Agilent 4200 TapeStation system. Paired-end sequencing (2×150 bp) was conducted on an Illumina NovaSeq6000 S4 flow cell. We obtained an average sequencing depth of 31,823,102 paired-end reads per sample. One sample producing low reads was excluded from subsequent analysis.

### Metagenomic processing, taxonomic profiling, and functional characterization

Paired-end libraries were demultiplexed using bcl2fastq2 v2.20.0.422 (Illumina), allowing up to one mismatch in the index barcode sequence (--barcode-mismatches 1). The resulting raw FASTQ reads were quality trimmed with Trimmomatic v0.36 and processed with fastp^115^ in order to remove poly-G tails. After quality trimming, reads were assessed for quality using FastQC and a consolidated report was produced using MultiQC^116^. High quality reads were mapped to the human genome (hg38) with metaWRAP^117^ to remove human derived sequences.

Taxonomic profiling was conducted using MetaPhlAn v4.0.4^33^ to generate relative abundances of microbial species identified in each sample. Functional characterization of the metagenomic reads was performed using HUMAnN 3.5^118^ using the ChocoPhlAn and UniRef90 databases (release 201901b). Reads per gene families were normalized for alignment quality and length using copies per million (CPM) units.

### Metagenomic assembly and functional analyses

MEGAHIT v1.2.9^119^ was used for independent de novo assembly of metagenomic samples and assembly quality was assessed with QUAST v5.0.2^120^. Genome bins were initially constructed from resulting scaffolds with >1,000 base pairs using three binning tools; CONCOCT^121^, MaxBin2^122^ and metaBAT2^123^. Subsequently, bins were refined and reassembled into metagenomically assembled genomes (MAGs) with metaWRAP^117^. CheckM v1.2.2^70^ used to assess the quality of MAGs by estimating several criteria established by MIMAG^69^, including completion, contamination, and presence of various rRNA and tRNA genes. Final MAGs fulfilling the criteria of medium-quality (≥50% completeness and ≤10% contamination) bins and other MIMAG standards were used for subsequent analyses. Taxonomic classification of all MAGs was assigned using GTDB-Tk v2.4.1 (release R226)^71^. Open reading frame prediction and functional annotation of microbial genes was performed with Bakta v1.6.1^124^, which resulted in 2,447,523 complete genes.

MAGs were dereplicated using FastANI v1.32^125^ clustering with a cutoff of ≥95% Average Nucleotide Identity (ANI), resulting in 221 unique MAGs. A total of 74 dereplicated MAGs showed no species-level classification in GTDB-Tk and no clustered representatives and were considered novel species. The 74 novel MAGs along with reference members of the same taxa in GTDB-Tk, were selected to generate alignments of their single-copy gene (SCGs) using hidden Markov model (HMM) profiles on GToTree v1.7.06^126^. Phylogenomic trees were inferred with FastTree v2.1.11^127^ and visualized on iToL^128^.

To better understand the metabolic capacity of the novel MAGs, DRAM v1.4.6 (Shaffer et al., 2020) was used to curate gene annotations and infer functional categories. Pangenomics analysis and identification of enriched functional pathways within the 221 dereplicated MAGs were performed using Anvi’o^129^ by annotating all protein coding genes using Clusters of Orthologous Genes release 2020^74^ and the Kyoto Encyclopedia of Genes and Genomes (KEGG) modules and classes^73^.

### Untargeted metabolomics using LC-MS and LC-MS/MS

#### Sample Preparation

A volume of 1 mL from the mouthwash samples (n=192) was added to an equal volume of acetonitrile (1:1 sample:acetonitrile). The sample were ultrasonicated for 5 minutes, centrifuged at 13,000 *g*, 4°C for 10 minutes and the supernatant (∼800 µL) was collected and dried under nitrogen flow (RapidVap N2, Labconco, Kansas City, MO, USA). The dried extract of the samples were reconstituted in 100 µL of 1:1 methanol:water. Blank samples (n=9) were included and treated in an identical manner as the samples. The reconstituted samples were stored at −80°C until analyzed. A pooled quality control (QC) sample was prepared by pooling equal volumes (20 µL) from each sample in the study, vortexed for 30 s and centrifuged at 13,000 *g*, 4⁰C for five minutes.

#### Analytical method

Untargeted metabolomics was employed using LC-MS to investigate the microbial metabolic profiles in the oral microbiome samples. Chromatography was performed using Vanquish UHPLC system (Thermo Fisher Scientific, Waltham, MA, USA) on a Zorbax RRHD HILIC Plus column (2.1 × 100 mm, 1.8 μm particle size, 95 Å, Agilent, Santa Clara, CA, USA) maintained at 40 °C. Mobile phases used were: (A) 0.1% formic acid, 10 mM ammonium formate in water and (B) 0.1% formic acid in acetonitrile. The gradient used was as follows: 0-10.5 min (5 to 60% A, 200 µL/min), 10.5-15 min (at 60% A, 200 µL/min), 15-17 min (60 to 5% A, 200 µL/min), 17-18 min (5% A, 200 to 300 µL/min), 18-31.50 min (to equilibrate at 5% A, 300 µL/min) and 31.5-32 min (5% A, 300 to 200 µL/min). The injection volume was 5 µL and the samples were maintained at 4°C during the analysis. A Tribrid Orbitrap Mass Spectrometer (Fusion Lumos, Thermo Fisher Scientific, Waltham, MA, USA) was used in switching ESI+ and ESI-modes for full LC-MS profiling and to generate data dependent MS/MS (ddMS/MS) accurate mass spectra for identification (most intense peaks). The operational parameters for both ESI modes were: spray voltage 3.5 kV, sheath, auxiliary and sweep gas flow rate were: 45, 8 and 1 (arbitrary units), respectively. Ion Transfer Tube and Vaporizer temperatures were maintained at 350 ⁰C and 300 ⁰C, respectively. Data were acquired in full scan mode with resolution 60,000 from *m/z* 50-1000. The ddMS/MS was performed at a resolution of 15,000 and stepped-normalized collision energies of 20, 30 and 50 at different mass range: *m/z* 50-250, *m/z* 250-500, *m/z* 500-750 and *m/z* 750-1000.

#### Metabolomics Analysis

The extracted samples were split into three different batches; each batch was randomized and analyzed in a single LC-MS analytical run. The pooled QC samples (n=6) were analyzed at the beginning of each batch to equilibrate the column prior analysis. The pooled QC sample injections were interspaced throughout the run to check the stability, robustness, repeatability, and performance of the analytical system. The QC sample was also extensively analyzed (n > 6) with ddMS/MS to include different mass ranges, a targeted list of in-house database standards, a list of expected metabolites, and a list of metabolite peaks with no fragmentation. Data pre-processing for untargeted peak-picking, alignment, and deconvolution, and metabolite annotation was carried out using Compound Discoverer v3.3 (Thermo Fisher Scientific, USA). The confidence in metabolite annotation was assigned as Level 1-4 (L1-L4) based on the recommendation by Chemical Analysis Working Group, Metabolomics Standards Initiative (MSI)^130^

### Statistical analyses

#### Oral microbial diversity analyses using 16S rRNA amplicons

Alpha diversity was assessed using species richness and Shannon’s Diversity Index, with reads rarefied to various depths between 1,000 and 10,000. For each depth, 100 iterations were performed and the mean values were used as the diversity estimates for each sample. Rarefaction curves indicated that 4,000 reads were sufficient to capture most of the bacterial diversity in this dataset. Samples with fewer than 4,000 reads were excluded, resulting in a final dataset with a total of 5,606,877 reads (63% of initial reads) and 380 ASVs from 628 samples. Comparison of alpha diversity metrics were performed by rarifying all samples to 4,000 reads. Kruskal–Wallis test was used to assess the differences in species richness between groups, with Dunn’s post-hoc applied for pairwise comparisons. A generalized linear mixed-effects model was used to identify covariates associated with Shannon Diversity Index, treating samples as a random effect.

#### Random forests classifier

We used a random forest classifier using the randomForest package in R to differentiate between obese and control groups using ASVs as predictors. The data was partitioned into training (80%) and testing (20%) sets, the model was fit using ten-fold cross validation repeated three times using 500 trees. The performance of the classifier was assessed by generating area under the receiver operating characteristic curves (AUC) using ROCR package^74^. The top features distinguishing between the two groups were determined using the varImp function in the caret package.

#### Oral microbial composition

For the amplicon sequencing, beta diversity was assessed at the ASV level using unweighted UniFrac, weighted UniFrac, and Bray–Curtis distances calculated by log+1 transformation of non-rarefied 16S rRNA gene count data. Unsupervised clustering was performed with Principal Coordinate Analyses (PCoA) using the phyloseq package and visualized with the ggplot2 package in R. For the metagenomics data, microbial composition was assessed by PCoA performed using Bray–Curtis distances using the species level relative abundances obtained from MetaPhlAn. PERMANOVA and EnvFit were performed with 10,000 randomizations using the *vegan* package. Covariates with *P* < 0.05 were considered statistically significant. Generalized linear mixed effect models with samples as the random effect were used to identify covariates associated with the top 3 Principal Coordinate (PCo) axes.

#### Oral microbiome-obesity associations

We first performed an association analysis using a full model that included 312 individuals, 27 non-collinear variables (VIF < 10), including 13 demography and lifestyle-related and 14 health-associated factors. This approach accounted for missing data while minimizing model overfitting, enabling us to evaluate the relationship between the oral microbiome and systemic health while adjusting for lifestyle-related factors. Association analyses were performed using analysis of the weighted UniFrac distances using two complementary approaches: PERMANOVA, which assesses microbiome distance-based differences between individuals, and EnvFit, which evaluates the top three principal coordinate dimensions explaining the greatest variation in the oral microbiome. To ensure the robustness of our findings, we repeated these analyses using a reduced model that included only the significant variables from the full model while incorporating the larger cohort of 618 individuals with available data. Across all these analyses, BMI emerged as a consistent factor associated with oral microbiome variation, highlighting it as a key variable for further investigation.

#### Differential abundance

MaAsLin2^34^ was used to identify ASVs, species, microbial pathways and enzymes that were differentially abundant between the obese and control groups. Multiple testing corrections were performed by computing the False Discovery Rates (FDRs) using the Benjamini–Hochberg method. Features with FDR-adjusted *P*< 0.05 were considered significant.

#### Metabolite abundances

Raw LC-MS peaks were subjected to batch-to-batch correction and total sum normalization. The dataset was then log transformed to restore normality. The metabolite (tR, m/z) pairs were exported with their normalized abundances for multivariate analysis (MVA) using Simca P16 (Sartorius Stedim Data Analytics AB, Sweden). The imported dataset of the samples in the study was mean-centered and unit-variance scaled. Principal component analysis (PCA), partial least squares - discriminant analysis (PLS-DA) and orthogonal partial least squares - discriminant analysis (OPLS-DA) were used for checking the analytical performance and modeling the differences between samples. Variable importance in the projection (VIP)>1.0 of the OPLS-DA, correlation scaled loading absolute p(corr) value>0.4, and a Student *t*-Test *p*-value<0.05 adjusted with false discovery rate using the Benjamini-Hochberg approach were used to identify the significantly altered metabolites in the study. Metabolite enrichment for structural classes, tissues, and dietary factors was performed using MetaboAnalyst and FDR-adjusted p-values<0.05 were considered significant^131^.

#### Procrustes analyses

To assess the relationship between microbial functional pathways and metabolite profiles, we performed Procrustes analysis, which evaluates the correlation between the distance matrices of two multivariate datasets by aligning their principal coordinate spaces and measuring structural similarity. We first constructed Bray-Curtis dissimilarity matrices for microbial functional pathways and metabolite profiles. Each dataset underwent Principal Coordinate Analysis (PCoA), and all available axes were used in Procrustes analysis, implemented in the vegan package in R. Procrustes analysis optimally rotates, scales, and translates one ordination space onto the other to maximize alignment, which was quantified using the Procrustes correlation coefficient (rho). To assess statistical significance, we conducted a permutation test with 1,000 permutations, randomly shuffling microbial and metabolite data points and recalculating the correlation each time. This was repeated to assess the relationship between microbial enzymes and metabolite profiles. The resulting P-values=0.0001 indicate that the observed correlation is highly unlikely to have occurred by chance, supporting a strong association between microbial enzymes, functional pathways, and metabolic composition.

#### Microbiome blood marker associations

We computed pairwise Spearman’s correlations between 366 bacterial species, 321 pathways, 285 metabolites, and 51 clinical markers. Multiple testing correction was performed by computing FDRs using the Benjamini-Hochberg method and FDR-adjusted *P*<0.05 were considered significant. Hierarchical clustering was applied to the correlation matrix to identify feature clusters. Enrichment of obesity related microbiome and clinical features in the clusters was evaluated with hypergeometric tests. An igraph object was created from the correlation matrix and global (degree centrality and betweenness) and cluster-specific (edge density and transitivity) network metrics were calculated. Central nodes in the network were inferred from global and cluster-specific metrics.

## Acknowledgments

This study is based upon work supported by Tamkeen under the Research Institute, New York University Abu Dhabi (NYUAD) (Grant No: G1206). We express our sincere gratitude to all volunteer participants, members of the Public Health Research Center (PHRC), and the Core Technology Platform Sequencing Center at NYUAD. We would also like to acknowledge the NYUAD High-Performance Computing Core team for providing computational resources. We are grateful to Jayaram Radhakrishnan in the NYUAD Bioinformatics Core for assistance with data preprocessing and bioinformatics and the members of Genetic Heritage Group for helpful discussions and constructive critique throughout this study.

## Author contributions

Conceptualization: A.R.J, Sample collection: A.A., R.A, Y.I., and The UAE Healthy Future Study Investigators Group, Experiments: T.W.D., C.E.L., L.U., G.Z., M.A, M.A. Data analysis: A.A.S., T.W.D., A.R.A., S.A, S.A.A, Y. M., M.A., N.D., and A.R.J. Interpretation: A.A.S., T.W.D., A.R.A., S.A, S.A.A, Y.I., and A.R.J. Original draft: A.A.S., T.W.D., A.R.A., S.A, and A.R.J. Writing and editing subsequent versions: A.A.S., T.W.D., A.R.A., S.A, S.A.A, Y.I., and A.R.J. Supervision: A.R.J, S.A.A., Y.I., R.A. Funding: A.R.J, Y.I., R.A. All authors discussed the results, contributed to the final manuscript, and have read and approved the submitted version.

## The UAE Healthy Future Study Investigators Group

Scott Sherman, Muna Tahlak, May Raouf, Mahera Abdul-Rahman, Hamda Khansaheb, Abdullah Al Naeemi, Fayza Al Ameri, Abdullah Al Junaibi, Eiman Al Zaabi, Naima Oumeziane, Asma Al Nuaimi, Marina Bastaki, Huda Al Shamsi, Hiba Al Humaidan, Mohammed Al Houqani, Fatma Al Maskari, Juma Al Kaabi, Ayesha Al Dhaheri, Syed Shah, Luai Ahmed, Fatme Al Anouti, Tom Loney, Alawi al Sheikh Ali, Mohammed Hag-Ali, Habiba Al Safar, Abderrahim Oulhaj, Wael Al Mahmeed, Mahmoud Traina, Khalid Saleh, Asma Al Mannaei, Omniyat Al Hajeri, Erik Koornneef, Laila Abdel Wareth, Mai Ahmed Sultan Essa Aljaber, Basema Saddik, Ghuwaya Al Neyadi, Walid Zaher, Hayfa Hamed, Buthaina Abdulla, Jamila Yacoub, Ali Abdul Kareem Al Obaidli, Andrea Jabari, Amar Ahmad, Yvonne Valles, Fatima Al Maisary, Imane Morjane, Sara Al Balushi, Mitha Albalushi, Manal Taimah Kayed, Vinu Manikandan, Manal Al Balooshi, Ayesha Al Hosani, Khaloud Alremeithi, Thekra Al Zaabi, Tamadher Alameri, Maryam Abed Al Marri, Mariem Elhadj, Fatma Al Shuhoomi, Ghausia Begum, Klaithem Mohamed, Manar T. Al Shaikh, Hassan Sabour Abdrabo, and Judalyn Del Monte.

